# Change in sexual signaling traits outruns morphological divergence in a recent avian radiation across an ecological gradient

**DOI:** 10.1101/424770

**Authors:** Guillermo Friis, Borja Milá

**Affiliations:** National Museum of Natural Sciences, Spanish National Research Council (CSIC), Madrid 28006, Spain

**Keywords:** *Junco*, sexual signaling, plumage coloration, phenotypic divergence, speciation, avian radiation

## Abstract

The relative roles of natural and sexual selection in promoting evolutionary lineage divergence remains controversial and difficult to assess in natural systems. Local adaptation through natural selection is known to play a central role in adaptive radiations, yet secondary sexual traits can vary widely among species in recent radiations, suggesting that sexual selection may also be important in the early stages of speciation. Here we compare rates of divergence in ecologically relevant traits (morphology) and sexually selected signaling traits (coloration) relative to neutral structure in genome-wide molecular markers, and examine patterns of variation in sexual dichromatism to understand the roles of natural and sexual selection in the diversification of the songbird genus *Junco* (Aves: Passerellidae). Juncos include divergent lineages in Central America and several dark-eyed junco (*J. hyemalis*) lineages that diversified recently as the group recolonized North America following the last glacial maximum (c.a. 18,000 years ago). We found an accelerated rate of divergence in sexually selected characters relative to ecologically relevant traits. Moreover, a synthetic index of sexual dichromatism comparable across lineages revealed a positive relationship between the degree of color divergence and the strength of sexual selection, especially when controlling for neutral genetic distance. We also found a positive correlation between dichromatism and latitude, which coincides with the latitudinal pattern of decreasing lineage age but also with a steep ecological gradient. Finally, we detected an association between outlier loci potentially under selection and both sexual dichromatism and latitude of breeding range. These results suggest that the joint effects of sexual and ecological selection have played a role in the junco radiation and can be important in the early stages of lineage formation.

## Introduction

Understanding the relative roles of natural and sexual selection in promoting evolutionary lineage divergence and speciation remains a central question in evolutionary biology, yet a challenging one to address in natural systems. Sexual selection has long been considered a significant driver of evolutionary diversification and speciation (Darwin 1871; Lande 1981; West-Eberhard 1983; Barraclough et al. 1995; Panhuis et al. 2001). However, the specific role of sexual selection in promoting phenotypic differentiation and lineage divergence remains controversial (Ritchie 2007b; Seddon et al. 2008; Kraaijeveld et al. 2011; Seddon et al. 2013). A particular mechanism of speciation by sexual selection has been proposed to operate through the acceleration of the rate of phenotypic change, which may in turn promote differences among allopatric populations in sexually selected traits involved in mate recognition (Price 1998; Seddon et al. 2013; Rowe et al. 2015). This process can lead to fast phenotypic differentiation (Panhuis et al. 2001), and might be especially relevant in the early stages of the speciation process (Ritchie 2007b; Seddon et al. 2008; Kraaijeveld et al. 2011). Indeed, several cases of highly variable secondary sexual traits in recently radiated systems have been documented, suggesting that sexual selection may account for part of the variation among closely related species of spiders (Masta and Maddison 2002), frogs (Boul et al. 2007), electric fishes (Arnegard et al. 2010) and birds (Young et al. 1994; Seddon et al. 2013; Safran et al. 2016; Wilkins et al. 2016).

Rapid divergence among isolated populations driven by sexual selection can be caused initially by random changes (drift) in sexually selected traits and the coevolution of correlated mate preferences, leading to differences in ornamental traits and mating success through so-called ‘runaway selection’ (Fisher 1930; West-Eberhard 1983; Questiau 1999). However, sexual signals necessarily interact with the environmental background and evolve in an ecological context, so that population divergence may be the result of the combined effects of sexual and natural selection (van Doorn et al. 2009; Maan and Seehausen 2011; Butlin et al. 2012; Seehausen et al. 2014). The combination of ecological opportunity and sexual selection has been invoked to explain lineage formation in the early stages of speciation and in recent adaptive radiations (e.g. Wagner et al. 2012; Scordato et al. 2014). Correlations of sexual selection with ecological parameters like latitude, habitat type, or migratory behavior have also been reported (Fitzpatrick 1994; Price 1998; Friedman et al. 2009; for review see Badyaev and Hill 2003) lending support to the hypothesis of sexual and ecological factors jointly driving lineage divergence. However, our understanding of the complex interactions and relative contributions of sexual and natural selection to the diversification process is still limited (Maan and Seehausen 2011; Safran et al. 2016).

Studies of sexually and ecologically selected traits in recent radiations that include lineages of different ages are particularly useful for gaining insight into the relative roles of sexual and ecological selection in driving lineage differentiation (Badyaev and Hill 2003; Kraaijeveld et al. 2011). Comparing the degree of divergence in ecomorphological and sexually selected traits allows assessing their rates of phenotypic change and thereby, the relative contributions of sexual and ecological pressures to the diversification process (Ritchie 2007a; Arnegard et al. 2010; Safran et al. 2013; Martin and Mendelson 2014). Biological systems presenting different spatial settings and occupying distinct environments also allows studying the evolution of sexual selection in relation to the demographic history or the colonization of new habitats (Endler 1980; Price et al. 2008; Wagner et al. 2012). Furthermore, by studying the evolution of sexual dichromatism, a common proxy of the intensity of sexual selection in avian taxa (Owens and Hartley 1998; Dunn et al. 2001; Huang and Rabosky 2014; Cooney et al. 2017) it is possible to test potential correlations between the rate of phenotypic divergence and the strength of sexual selection across different lineages.

Systems encompassing both old and recently radiated lineages showing variation in ecological and sexually selected traits may be found in taxa that underwent range expansions and colonized new areas across latitudinal gradients following glacial periods (Schluter 2000; Coyne and Orr 2004). Ecological adaptations in some of these systems are accompanied by latitudinal variation in potential sexually selected traits (e.g. New World warblers, orioles, Hamilton 1961), suggesting concomitant effects of natural and sexual selection. One such system is the songbird genus *Junco*, a species complex that includes highly divergent phylogenetic lineages in Central America as well as recently diversified lineages in temperate North America. Previous molecular studies indicate that northern juncos represent a case of recent radiation from a Central American ancestor during the recolonization of North America after the last glacial maximum (LGM), c.a. 18,000 years ago (Milá et al. 2007; Friis et al. 2016). The Central American taxa to the south of the distribution include the divergent volcano junco (*Junco vulcani*) in Costa Rica; Baird’s junco (*Junco bairdi*) from the southern tip of the Baja California Peninsula; the island junco (*Junco insularis*) on Guadalupe Island in the Mexican Pacific; and two closely related yellow-eyed juncos in the highlands of Chiapas (Mexico) and Guatemala, currently classified as *Junco phaeonotus fulvescens* and *Junco phaeonotus alticola*, respectively. Post-glacially radiated lineages across the North American continent comprise two more yellow-eyed taxa in mainland Mexico, *Junco ph. phaeonotus* and *Junco ph. palliatus*, and at least six forms currently grouped within the dark-eyed junco (*Junco hyemalis*) complex: the red-backed junco (*J. h. dorsalis*) from southwestern USA; the gray-headed junco (*J. h. caniceps)* in the Rocky Mountains; the Oregon junco (*J. h. oreganus*) across the West, composed in turn of several distinct forms from northern Baja California to Alaska, including *townsendi*, *pontilis*, *thurberi*, *pinosus*, *montanus*, *shufeldti* and *oreganus*; the pink-sided junco (*J. h. mearnsi*) in the northern Rocky Mountains; the white-winged junco (*J. h. aikeni*) in the Black Hills of South Dakota; and the slate-colored junco in eastern and boreal North America, comprising *J. h. hyemalis*, *J. h. carolinensis* and *J. h. cismontanus* (Fig. 1A, Table 1; Miller 1941; Sullivan 1999; Nolan et al. 2002). The marked diversity of plumage patterns and colors among the recently radiated northern forms of junco (Fig. 1A) suggests that sexual selection may have played a relevant role in the phenotypic diversification of the young forms of junco. Nevertheless, the fact that the radiation took place across a wide latitudinal axis of pronounced ecological variability suggests potential interactions between sexual selection and ecological selective pressures related to northern habitats (e.g. Fitzpatrick 1994; Price 1998; Friedman et al. 2009; for review see Badyaev and Hill 2003).

**Figure 1.**
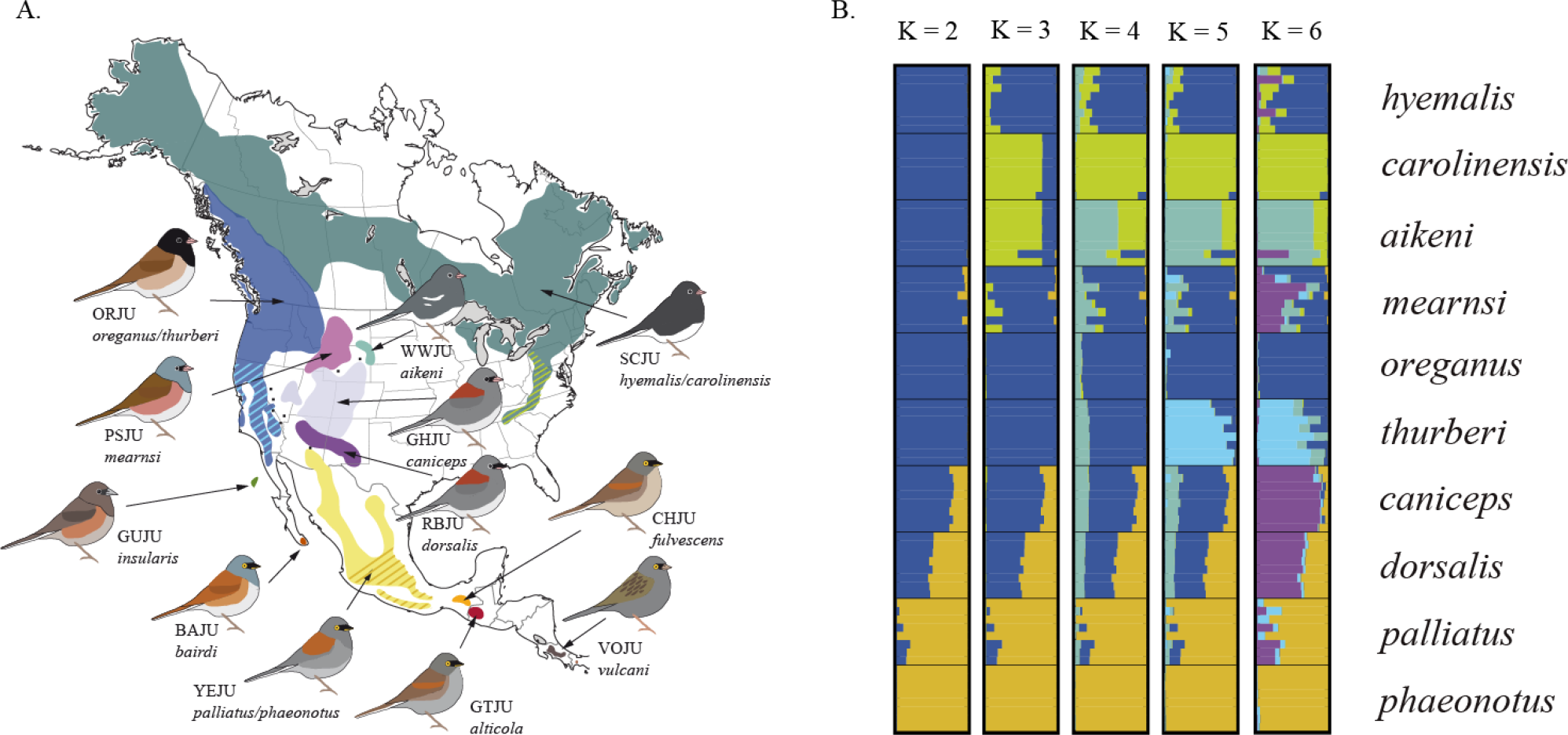
Geographic distribution of phenotypic variation in the genus *Junco* and neutral genetic structure of the northern forms. (A) Distribution map of the different junco forms. Colored areas correspond to the breeding ranges of the major forms (see Table 1 for a detailed nomenclature). Dots represent isolated localities with hybrid/intermediate individuals and the striped areas correspond to subspecific forms *carolinensis* (light green), *phaeonotus* (brown) and *thurberi* (light blue). (B) Genetic structure of the northern junco forms from a STRUCTURE analysis based on 11,698 selectively neutral genome-wide SNPs for K = [2-6]. Each horizontal bar corresponds to an individual, with different colors corresponding to posterior assignment probabilities to a given number (K) of genetic clusters. Colors correspond to those on the range map on Fig. 1A.

**Table 1.**
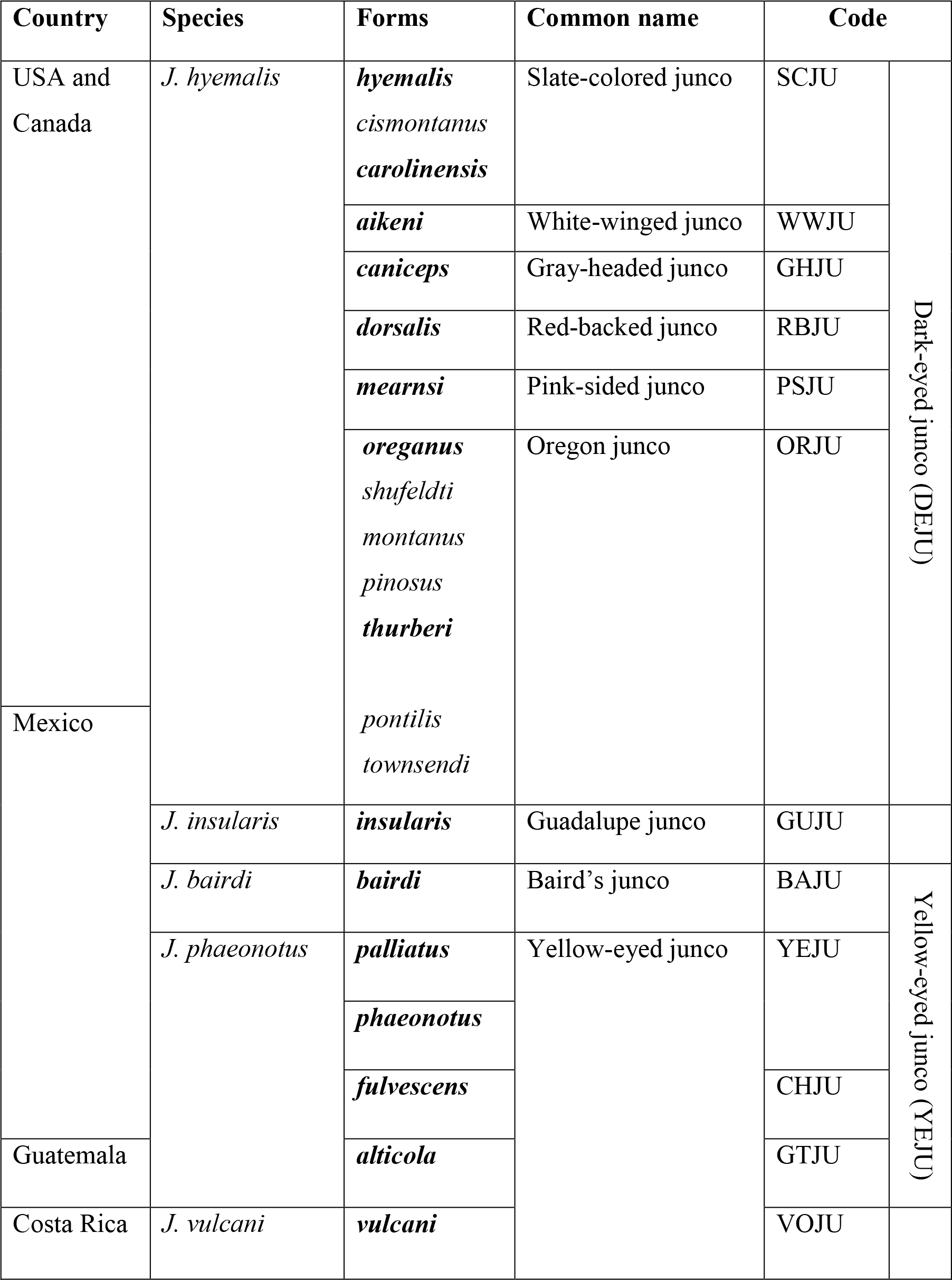
Taxonomy of junco forms. Those forms included in this study are shown in bold. The five species-level taxa are those currently recognized by the American Ornithologists’ Society (2017).

Here, we study patterns of genetic and phenotypic differentiation in the genus *Junco*, including older Central American species and recently radiated North American lineages, and infer the relative roles of sexual selection and ecological factors in driving diversification. We first study the general patterns of neutral genetic structure in the recently radiated northern junco lineages using genome-wide single nucleotide polymorphisms (SNPs) obtained with genotyping-by-sequencing (GBS, Elshire et al. 2011). Then we use morphometric and spectrophotometric data from museum specimens with three major aims: (i) comparing rates of phenotypic evolution in both traits of ecological importance and plumage coloration by means of discriminant function analyses (DFA) and Mahalanobis distances to assess the relative contributions of ecological and sexual selection; (ii) assessing potential interactions between divergent mate choice and selective ecological pressures by testing for positive correlation between latitude and sexual dichromatism across the distribution of the genus with multivariate and linear regression analyses; (iii) testing the role of sexual selection in driving diversification by examining the correlation between the degree of divergence on sexually selected characters and a synthetic index of sexual dichromatism by means of simple and partial Mantel tests; and (iv) applying redundancy analysis to explore the potential concomitant effects of ecological divergence and differential mate choice in shaping genetic adaptive variability by testing for associations between allele frequencies and both latitude and sexual dichromatism.

## Materials and methods

### Population sampling

Adult, territorial male juncos were sampled across their range using mist nets in order to obtain phenotypic data and blood samples for DNA extraction. Each captured individual was aged, sexed, and marked with a numbered aluminum band. A blood sample was collected by venipuncture of the sub-brachial vein and stored in Queen’s lysis buffer (Seutin 1991) or absolute ethanol at −80ºC in the laboratory. After processing, birds were released unharmed at the site of capture. All sampling activities were conducted in compliance with Animal Care and Use Program regulations at the University of California Los Angeles, and with state and federal scientific collecting permits in the USA and Mexico. A high-quality tissue sample for whole-genome sequencing was obtained from a slate-colored junco (*J. hyemalis carolinensis*). Genomic DNA was extracted from blood and tissue samples using a Qiagen DNeasy kit (Qiagen^TM^, Valencia, CA) for downstream analyses.

### Genotyping-by-sequencing

We used genotyping-by-sequencing (Elshire *et al*. 2011) to obtain individual genotypes from 243 juncos belonging to the following taxa (with sample sizes in parentheses): *hyemalis* (14), *carolinensis* (22), *aikeni* (12), *mearnsi* (12), *oreganus* (16), *thurberi* (34), *caniceps* (69), *dorsalis* (48), *palliatus* (8) and *phaeonotus* (8) (Table 2, Table S1 from Supplementary Information). GBS libraries were prepared and sequenced at Cornell University’s Institute for Genomic Diversity, using the restriction enzyme PstI for digestion. Sequencing of the 243 individually-barcoded libraries was carried out in five different lanes (along with other 232 junco samples intended for other analyses) of an Illumina HiSeq 2000, resulting in an average of 243.2 million good barcoded single-end reads 100 bp in length per lane.

**Table 2.**
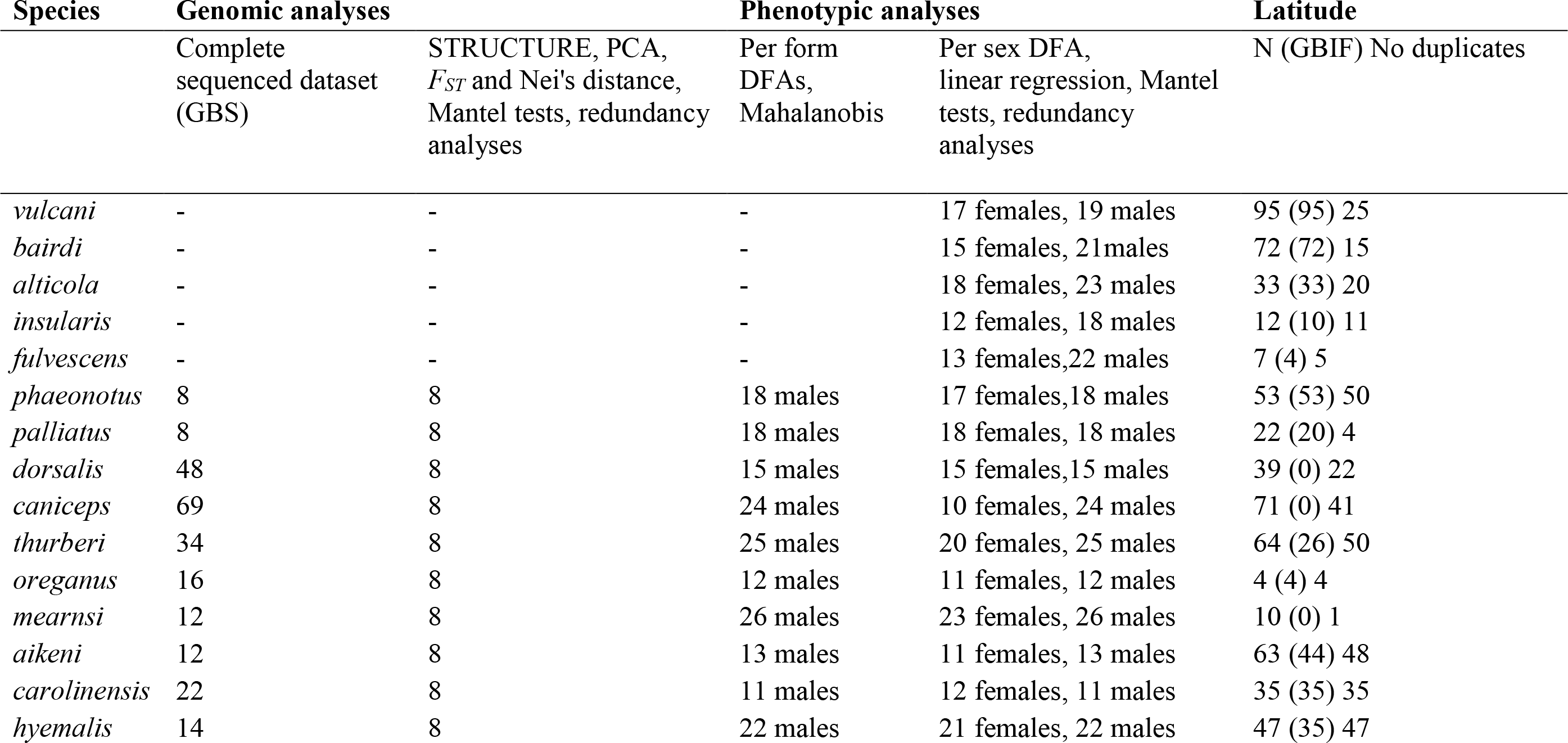
Sample sizes of the different datasets used in the study, including sequencing and genomic analyses, multivariate analyses on phenotypic data and collection of latitudinal records.

### Genome assembly, GBS reads alignment and variant calling

A high quality genome of *Junco hyemalis* sequenced and assembled by Dovetail™ by means of Hi-C (Belton et al. 2012) libraries based on Chromosome Conformation Capture (for details see Friis et al. *in press*) to be used as reference. To recover the chromosomal coordinates of the obtained scaffolds we mapped and oriented them against the zebra finch (*Taeniopygia guttata*) genome v87 available in Ensembl (Yates et al. 2016). We used the Chromosembler tool available in Satsuma (Grabherr et al. 2010) resulting in a final genome assembly of 955.9 Mb length and a N50 of 71.46 Mb.

Because Hi-C approach failed in sequencing the sexual chromosome Z, we recovered it from a draft consensus genome assembled by combining low-coverage genomes of eight different junco individuals intended for a parallel study (See Supplementary Information for details), once again using Chromosembler and the Z chromosome of the zebra finch. We evaluated GBS read quality using FASTQC (Andrews 2010) after sorting them by individual with AXE (Murray and Borevitz 2017) and performed the trimming and quality filtering treatment using Trim Galore (Krueger 2015), excluding all reads out of a range length between 40 and 90 bp long. Adapter removal stringency was set to 1 and the quality parameter ‘q’ to 20. GBS reads were then mapped using the mem algorithm in the Burrows-Wheeler Aligner (BWA, Li and Durbin 2009). Read groups assignment and BAM files generation was carried out with Picard Tools version 2 (http://broadinstitute.github.io/picard). We used the Genome Analysis Toolkit (GATK, McKenna et al. 2010) version 3.6-0 to call the individual genotypes with the HaplotypeCaller tool. We finally used the GenotypeGVCFs tool to gather all the per-sample GVCFs files generated in the previous step and produce a set of jointly-called SNPs and indels (GATK Best Practices, DePristo et al. 2011; Auwera et al. 2013) in the variant call format (*vcf*). Because GBS data does not provide enough coverage for base quality score recalibration, we used VCFTOOLS (Danecek et al. 2011) to implement a ‘hard filtering’ process, customized for each of the downstream analyses (see below).

### Genetic structure analyses

To explore genome-wide population structure among recently diverged junco forms, we ran a STRUCTURE (Pritchard et al. 2000) analysis based on SNP data. Using VCFTOOLS, we retained the eight samples of each population with the lower proportion of missing sites for a final number of 80 samples (Table 2). We constructed a data matrix of biallelic SNPs excluding those out of a range of coverage between 4 and 50 or with a genotyping phred quality score below 40. Positions with less than 75% of individuals genotyped for each taxon were removed from the data matrix, along with those presenting a minor allele frequency (MAF) below 0.02. We implemented a threshold for SNPs showing highly significant deviations from Hardy-Weinberg equilibrium (HWE) with a p-value of 10^−4^ to filter out false variants arisen by the alignment of paralogous loci. We used BayeScan (Foll and Gaggiotti 2008) to compute per SNP posterior probabilities of being under divergent or balancing selection in order to (i) filter them out for analysis of neutral genetic structure and (ii) study how adaptive variability is structured across the genome and test for potential correlations with proxies of selective forces (See Adaptive variation association tests section). BayeScan computes and decomposes per-SNP *F_ST_* scores into a population-specific component shared by all loci that approximates population related effects, as well as a locus-specific component shared by all populations, which accounts for selection. BayeScan compares two models of divergence, with and without selection, and assumes a departure from neutrality when the locus-specific component is necessary to explain a given diversity pattern (Foll 2012). We used BayeScan with default settings and a thinning interval size of 100 to ensure convergence. For each SNP we obtained the posterior probability for the selection model and the *F_ST_* coefficient averaged over populations. For outlier detection and exclusion, we implemented a false discovery rate of 0.1. To filter out the SNPs under linkage disequilibrium (LD) we used the function snpgdsLDpruning from the SNPrelate package (Zheng 2012) in R Studio (R_Studio_Team 2015) version 1.0.136 with R (R_Core_Team 2015) version 3.2.2. We applied the correlation coefficient method with a threshold of 0.2 (method ="corr", ld.threshold=0.2), resulting in a final data matrix of 11,698 SNPs. We converted the *vcf* file to STRUCTURE format using PGDspider (Lischer and Excoffier 2012) version 2.0.5.1. Bash scripts to perform the analyses were created with STRAUTO (Chhatre and Emerson 2016) and we ran the program five times per K, for values of K ranging from 1 to 10 after running a preliminary analysis to infer the lambda value. The burn-in was set to 50K iterations and the analysis ran for an additional 100K iterations. Similarity scores among runs and graphics were computed with CLUMPAK (Kopelman et al. 2015).

We used the same SNP data matrix to examine population structure by means of a principal components analysis (PCA). We used the function snpgdsPCA available in SNPrelate to perform the PCA and obtain the eigenvectors to be plotted. Finally, we computed a matrix of pairwise Nei’s distances and *F_ST_* values from the same SNP dataset used for the PCA and the STRUCTURE analysis using the R packages adegenet (Jombart 2008) and hierfstat (Goudet et al. 2015), respectively.

### Morphometric data and divergence analysis

We obtained morphometric data from 531 museum specimens representing all main junco forms, deposited at various natural history museums (see Table 2, Appendix I from Supplementary Information). A wing ruler was used to measure unflattened wing length to the nearest 0.5 mm, and dial calipers of 0.1-mm precision were used to measure tail length, tarsus length, exposed bill culmen, and bill width and depth. All measurements were taken by a single observer (BM) following Milá *et al*. (2008). We examined overall morphological differentiation among northern junco forms (*phaeonotus*, *palliatus*, *dorsalis*, *caniceps*, *thurberi*, *oreganus*, *mearnsi*, *aikeni*, *carolinensis* and *hyemalis*) using male data in a discriminant function analysis (DFA) after transforming all variables using natural logarithms. Analyses were conducted in R Studio 1.0.136 with R 3.2.2.

### Colorimetric data and divergence analysis

We obtained colorimetric data from the same 531 museum specimens measured for morphometric analysis (Table 2, Appendix I from Supplementary Information). To collect reflectance spectra we used a JAZ-EL200 spectrophotometer with a deuterium-tungsten light source via a bifurcate optical fiber probe (Ocean Optics^TM^). The reflectance captor probe was mounted on a black rubber holder which excluded all external light and maintained the probe fixed at a distance of 3 mm from the feather surface at a 90° angle (e.g. Schmitz-Ornes 2006; Chui and Doucet 2009). The spectrum of each measurement ranged from 300 to 700 nm and consisted of three replicate measurements of three different readings per replicate, taken on each of six plumage patches: crown, nape, back, breast, flank and belly. Replicates were averaged before analysis. All reflectance data is expressed as the percentage of reflectance from a white standard (WS-1, Ocean OpticsTM). The white standard was measured after each specimen and the spectrophotometer was recalibrated regularly. All measurements were taken by a single observer (GF).

We obtained colorimetric variables by applying the avian visual model by Stoddard and Prum (2008), based on Goldsmith’s (1990) tetrahedral color space for spectral data. We used the R-package pavo (Maia et al. 2013a) to calculate the relative quantum catch for each cone using the function vismodel. Specifically, we applied the visual system, sensitivity and ocular environmental transmission of the blue tit as available in the package, the ‘forestshade’ illuminant option and an ideal homogeneous illuminance for the background. We also applied the von Kries color correction transformation. We then obtained the spherical coordinates of tetrahedral color space describing the hue (Θ and φ) and the achieved chroma (r_a_) using the function colspace. We included the normalized brilliance as a fourth variable, computed as described in Stoddard and Prum (2008). Once we had computed the avian visual model variables, we used the R function boxplots.stats to detect and exclude eleven potentially wrongly measured samples be implementing a highly conservative coefficient of 10, *i.e*. those data measures 10 times higher or lower than the length of the third and fourth interquartile range. Once again, we applied DFA to resulting dataset to examine overall patterns of color differentiation among northern junco forms (*phaeonotus*, *palliatus*, *dorsalis*, *caniceps*, *thurberi*, *oreganus*, *mearnsi*, *aikeni*, *carolinensis* and *hyemalis*) using male data.

### Rates of trait divergence analysis

In order to compare rates of phenotypic divergence between sets of ecomorphological and secondary sexual traits in northern, recently diversified juncos (*phaeonotus*, *palliatus*, *dorsalis*, *caniceps*, *thurberi*, *oreganus*, *mearnsi*, *aikeni*, *carolinensis* and *hyemalis*), we computed pairwise Mahalanobis distances (Mahalanobis 1936), a measure of dissimilarity scaled by the variation within groups and applicable to multivariate trait spaces (e.g. Eldredge et al. 2005; Arnegard et al. 2010). We used the pairwise.mahalanobis function from the HDMD v1.2 R package and computed the square root of the resulting value to obtain the pairwise distances for morphological and colorimetric variables separately. We also ran a linear regression analysis between pairwise values of trait distance and Nei’s genetic distance to study differences in correlation patterns of both set of phenotypic values with neutral genetic differentiation (Arnegard et al. 2010).

### Differential sexual selection analysis

In order to estimate differential intensity of sexual selection among all junco lineages, we computed a synthetic index of the overall differences between females and males to compare the degree of dichromatism among lineages applying multivariate analysis. To calculate this index, we first performed a DFA by sex. Because comparisons among scores of different multivariate analysis and datasets are not statistically valid, we did not separate the analysis for different forms, and ran the DFA for the entire sample space (Montgomerie 2006). Second, we computed the DFA score means of females and males of each form and their 95% confidence intervals (CI). Third, for each one of these per junco form values, we subtracted the average of male and female DFA scores resulting in zero-centered differences between sexes, for clearer graphic comparison.

We also conducted a linear regression between the degree of averaged dichromatism and mean geographical coordinates of each form along the latitudinal axis of the distribution of the juncos. To compute the latitudinal means, we used the geographic locations of our own field sampling, complemented with GBIF accessions for each junco form (Table 2, Table S2).

Finally, to test the relationship between sexual selection and plumage color diversification, we used the R package vegan (Oksanen et al. 2016) to run a simple Mantel test (Mantel 1967) between pairwise Mahalanobis distances based on color variables and the pairwise sum of the scores of the sexual dichromatism index as an estimate of the intensity of sexual selection experienced by the two lineages under comparison (Seddon et al. 2013). We also ran a partial Mantel test (Smouse et al. 1986) to control for neutral genetic divergence, including the matrix of pairwise Nei’s distances to be partialed out. Complementarily, we ran a second simple Mantel test to test for a correlation between the two independent matrices (sexual dichromatism and genetic distance). Significance was computed through 9,999 matrix permutations. Analyses were carried out in R version 3.2.2 and SPSS v22 (See Appendix II from Supplementary Information for R scripts).

### Adaptive variation association tests

We tested for associations of adaptive variation in the northern lineages with sexual dimorphism and latitude, as proxies of sexual selection and ecological selective pressures, respectively, using redundancy analysis (RDA, Van Den Wollenberg 1977; Legendre and Legendre 1998). Because of the high collinearity between latitude and sexual dichromatism (see Results), we ran RDAs separately for the two variables to obtain an ordination over a single explanatory variable (Lepš and Šmilauer 2003; Borcard et al. 2011) and then performed a variance partition test to assess the degree of overlapping between each variable’s explained variance. The response variable was the frequency of the less frequent allele for each one of the biallelic SNPs putatively under selection detected by BayeScan when using a FDR of 0.1 (Meirmans 2015; Rellstab et al. 2015), computed over each of the young northern junco forms (See Genetic structure analyses section for details of BayeScan analysis). The explanatory variables were averaged latitude and sexual dichromatism per form as previously described. We ran the redundancy analyses using the rda function available in the R package vegan and obtained their statistical significance by a permutation-based procedure with 9,999 permutations. The variance partition analysis was carried out with the varpart R function, also available in vegan (See Appendix II from Supplementary Information for R scripts).

## Results

### Neutral genetic structure among northern *Junco* forms

The STRUCTURE analysis of the young junco lineages for two genetic clusters (K = 2) showed a gradual pattern of divergence from the Mexican *J. p. phaeonotus* to the *J. h. hyemalis* of Canada, approximately separating the yellow-eyed from the dark-eyed forms, with *caniceps* and *dorsalis* forms showing intermediate assignment probabilities, in congruence with their geographic positions. The analysis for K = 3 revealed *carolinensis* and *aikeni* as an independent cluster, yet in K = 4 a*ikeni* presented intermediate probabilities of belonging to a fourth, separated genetic group. In the test for five clusters (K = 5), *thurberi* also appeared as a separated population with little shared variance with other forms. In the test corresponding to K = 6, individuals from red-backed forms *dorsalis* and especially *caniceps* presented high assignment probabilities to a sixth independent cluster (Fig. 1B).

The PCA yielded similar general patterns. A plot of PC1 (5.9% of explained variance) against PC3 (4.2% of explained variance against 4.7% of the PC2, but showing better cluster resolution) revealed *carolinensis* and *aikeni* as highly differentiated groups and clear clustering for all the dark-eyed junco forms. Separation between the *J. phaeonotus* forms was less pronounced, and appeared as close groups to *dorsalis*, the neighbor dark-eyed form from southern USA (Fig. 2A). Plot of second and fourth components revealed similar degrees of clustering among junco forms (Fig. S1 from Supporting Information).

**Figure 2.**
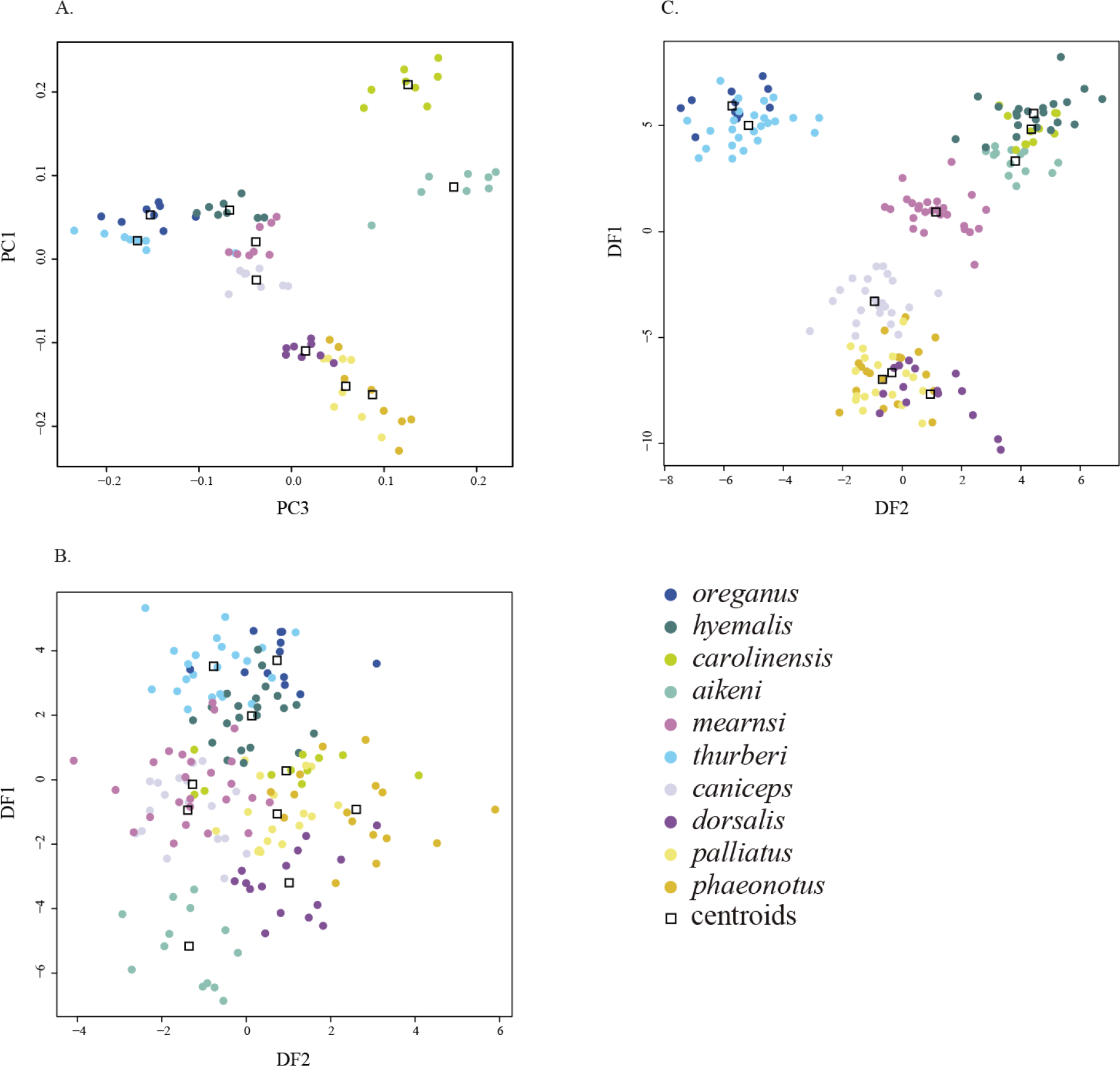
Neutral genetic structure and phenotypic differences among the recently radiated forms of junco. (A) Genetic structure of northern junco forms based on the first and third axis of a principal components analysis of selectively neutral genome-wide SNPs. (B) and (C) show the first two discriminant functions in a discriminant function analysis (DFA) based on morphological variables and plumage color variables, respectively. Marker colors correspond to those on the range map on Fig. 1A.

Nei’s distances and *F_ST_* values were generally congruent with genetic structure analyses. Southern forms *phaeonotus* and *palliatus* showed the highest values for both indices, while northern forms showed lower levels of pairwise differentiation with a clear increase in the *aikeni* and *carolinensis* forms (Table 3).

**Table 3.** Pairwise Nei’s genetic distances (lower diagonal) and *F_ST_* values (upper diagonal) for all northern junco forms based on 11,698 independent, selectively neutral SNP loci.

| Nei's\FST | *phaeonotus* | *palliatus* | *dorsalis* | *caniceps* | *thurberi* | *oreganus* | *mearnsi* | *aikeni* | *carolinensis* | *hyemalis* |
| --- | --- | --- | --- | --- | --- | --- | --- | --- | --- | --- |
| *phaeonotus* |  | 0.085 | 0.097 | 0.102 | 0.106 | 0.109 | 0.101 | 0.114 | 0.121 | 0.107 |
| *palliatus* | 0.048 |  | 0.082 | 0.087 | 0.095 | 0.095 | 0.089 | 0.102 | 0.108 | 0.094 |
| *dorsalis* | 0.056 | 0.051 |  | 0.071 | 0.086 | 0.082 | 0.077 | 0.089 | 0.092 | 0.080 |
| *caniceps* | 0.059 | 0.054 | 0.047 |  | 0.081 | 0.075 | 0.072 | 0.083 | 0.086 | 0.073 |
| *thurberi* | 0.062 | 0.058 | 0.055 | 0.051 |  | 0.081 | 0.078 | 0.091 | 0.093 | 0.079 |
| *oreganus* | 0.065 | 0.059 | 0.054 | 0.048 | 0.050 |  | 0.074 | 0.085 | 0.085 | 0.072 |
| *mearnsi* | 0.060 | 0.055 | 0.050 | 0.046 | 0.049 | 0.047 |  | 0.080 | 0.086 | 0.071 |
| *aikeni* | 0.068 | 0.064 | 0.059 | 0.055 | 0.057 | 0.055 | 0.052 |  | 0.094 | 0.083 |
| *carolinensis* | 0.074 | 0.069 | 0.062 | 0.057 | 0.059 | 0.056 | 0.056 | 0.062 |  | 0.081 |
| *hyemalis* | 0.063 | 0.058 | 0.052 | 0.047 | 0.049 | 0.045 | 0.045 | 0.053 | 0.052 |  |

### Patterns of morphometric differentiation

The plot of the first two discriminant functions from a DFA on morphometric variables for the northern lineages of *Junco* revealed a pattern of low clustering among groups (Fig. 2B). The forms *aikeni* and *dorsalis*, and to a lesser extent, *thurberi*, *oreganus* and *hyemalis* presented certain degree of separation. The *phaeonotus* centroid showed also a considerable divergence from other northern junco forms, but individuals showed considerable spread across multivariate space. The remaining forms presented extensive overlap.

### Patterns of color differentiation

In contrast to morphometric variables, the DFA based on spectral data revealed considerable differentiation in plumage coloration patterns. A plot for the first two discriminant functions showed clear separation of the two black-hooded Oregon junco forms, *oreganus* and *thurberi*, from the rest of lineages, as well as for *mearnsi* and *caniceps*, which occupied more centered positions. The two slate-colored forms, *hyemalis* and *carolinensis* clustered together with the phenotypically similar *aikeni*. Similarly, *phaeonotus* and *palliatus* showed no differentiation between them and overlapped with *dorsalis* (Fig. 2C). These patterns were remarkably congruent with the general neutral genetic structure recovered in the PCA (Fig. 2A) and with the geographic distribution of the northern juncos (Fig. 1A).

### Rates of trait divergence analysis

Pairwise Mahalanobis distances revealed high disparity in rates of divergence for morphological and colorimetric variables, ranging from 0.08 to 0.29 and from 2.87 to 26.94, respectively (Fig. 3A). Linear regression plots showed contrasting patterns of relative stasis in morphometric traits versus high evolvability in color traits in relation with genetic distances (Fig. 3B). The analysis yielded a highly significant correlation between color and genetic divergence (*P* = 1.04×10^−5^, R^2^ = 0.37), and was only marginally significant between morphometric and genetic distance (*P* = 0.11, R^2^ = 0.06).

**Figure 3.**
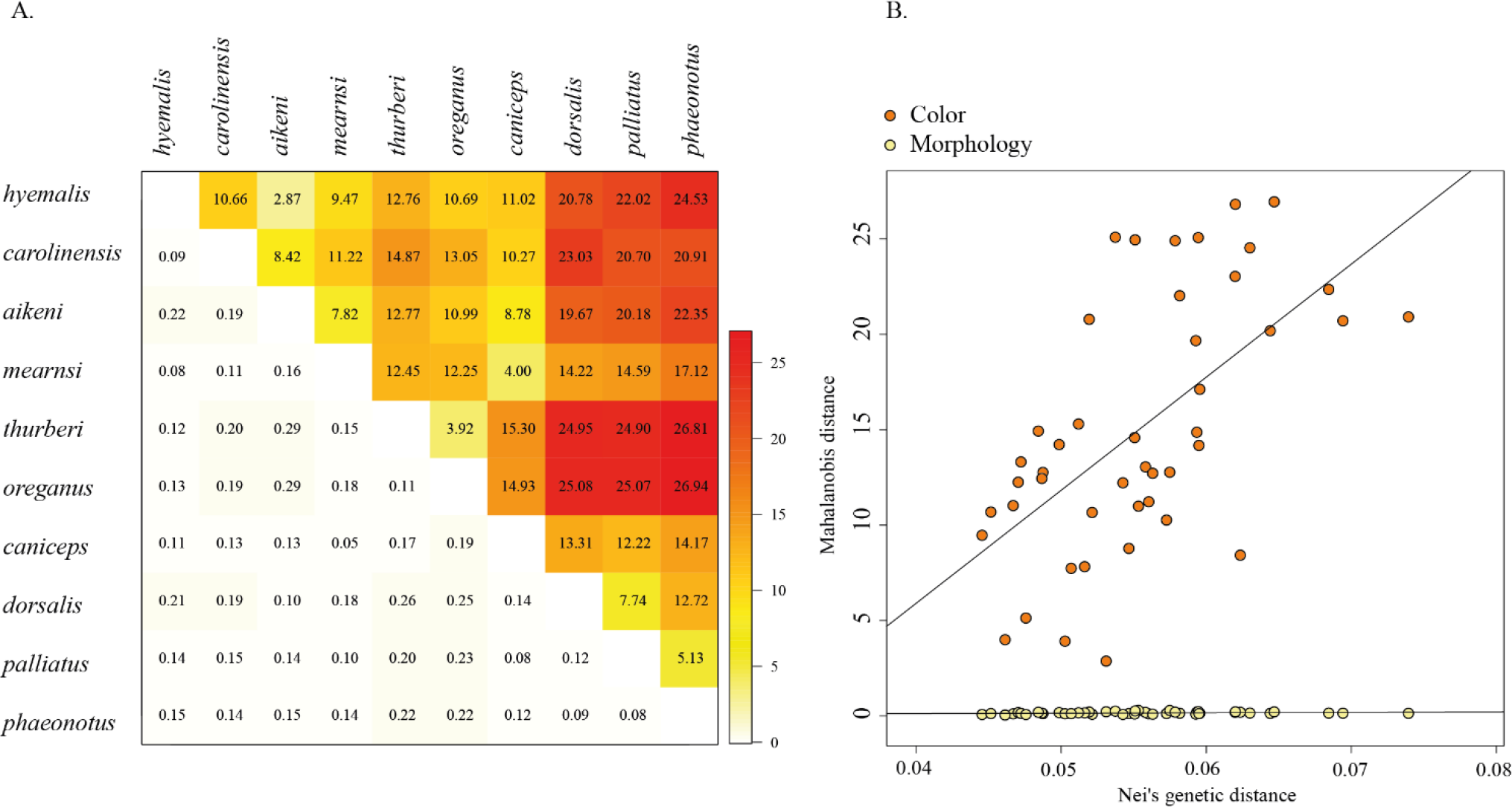
Phenotypic trait divergence rates and their correlation with genetic distance. (A) Pairwise Mahalanobis distances for colorimetric (above diagonal) and morphometric (below diagonal) traits for northern junco forms. (B) Linear regression of Nei’s genetic distance against pairwise Mahalanobis distances based on colorimetric traits (orange circles, *P* = 1.04×10^−5^, R^2^= 0.37) and morphometric traits (yellow circles, *P* = 0.11, R^2^ = 0.06).

### Differential sexual selection analysis

The sexual dichromatism index computed from the DFA scores revealed a gradually increasing pattern of differentiation between males and females when ordering the forms from south to north (Fig. 4A), with the exception of *caniceps* and *carolinensis*, which did not follow this pattern. The latitudinal signal of increasing dichromatism was also evident when considering only the recently radiated forms, where the yellow-eyed Mexican lineages presented the lowest male-female differentiation values in contrast to the most boreal forms, *hyemalis* and *oreganus*. The linear regression between mean male-female differences and latitude was highly significant (*P* = 4×10^−4^), with latitude explaining 63% of the variance in sexual dichromatism (R^2^ = 0.63). The Pearson correlation coefficient was equal to 0.79. Remarkably, the pattern persisted within the *oreganus* individuals of our study, with subspecies *thurberi* from northern Baja California showing lower dichromatism than northern *oreganus* from British Columbia (Fig. 4B). The simple Mantel test for color Mahalanobis distances and degree of sexual dichromatism revealed a moderate but significant correlation between the two measures (*P* = 0.048, r = 0.31). The correlation and significance increased when controlling for genetic distance in the partial Mantel test (*P* = 0.007, r = 0.42). In turn, we found a non-significant correlation between sexual dichromatism and Nei’s genetic distance (*P* = 0.85, r = −0.36).

**Figure 4.**
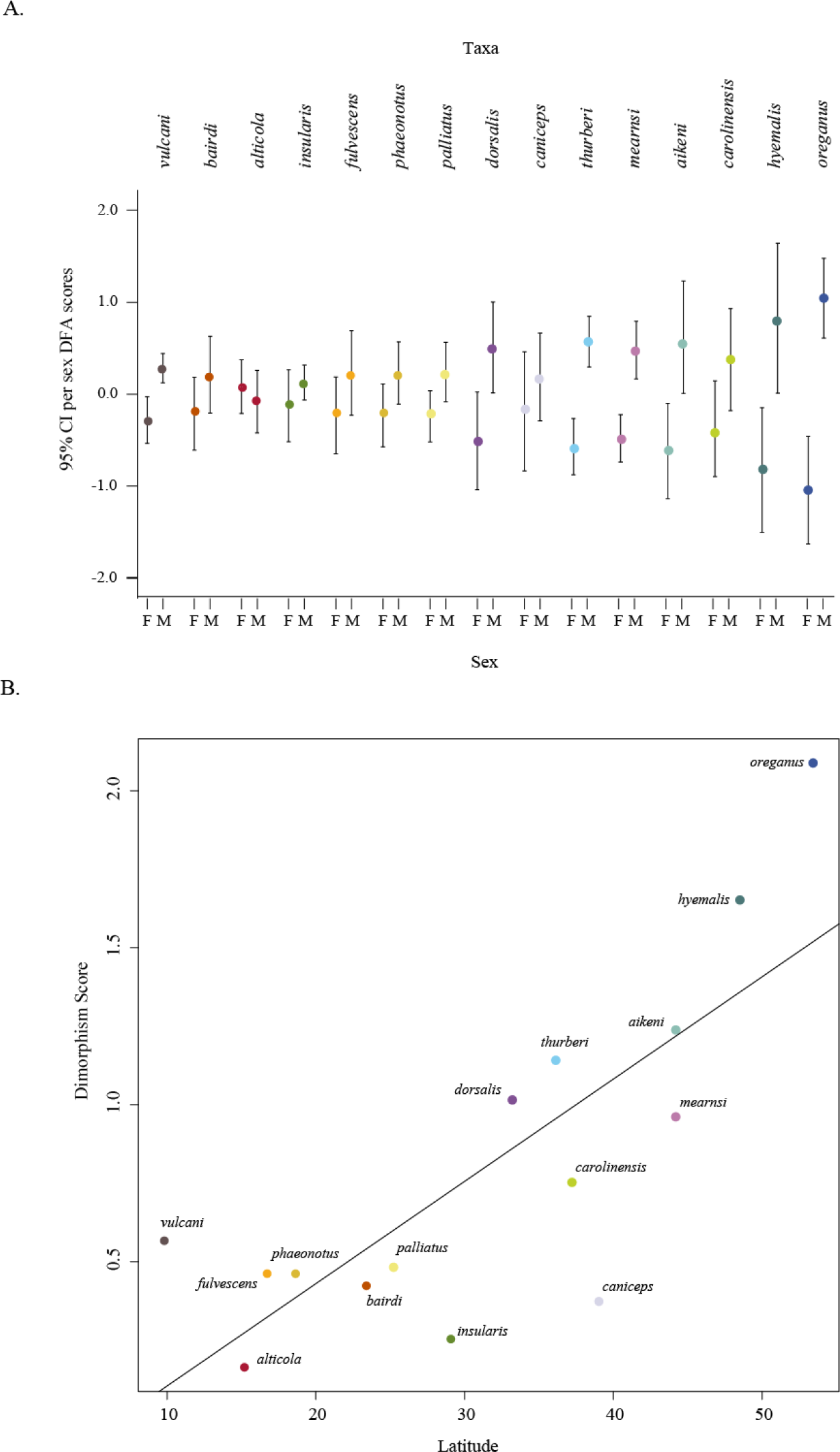
Latitudinal pattern of gradual increase in sexual dichromatism across the *Junco* distribution range. (A) Centered sex-discriminant DFA scores of avian visual model variables (Θ, φ, achieved chroma and normalized brilliance) across the entire sample space for junco forms ordered from south to north. (B) Lineal regression between the degree of average dichromatism and mean latitude for each form (*P* = 4×10^−4^, R^2^ = 0.63, Pearson correlation coefficient = 0.79).

### Adaptive variation association tests

BayeScan genomic survey yielded 113 outliers putatively under divergent selection distributed across the genome from an initial dataset of 24,792 SNPs when applying a FDR of 10%, four of them located in the Z chromosome (see the Manhattan plot of posterior probabilities in Fig. S2 in the Supplementary Information). Redundancy analyses revealed that both sexual dichromatism and latitude had significant effects on adaptive genomic variance, with *P* values equal to 0.02 and 0.002, respectively. Sexual dichromatism explained 23% of the total adaptive variance (adjusted R^2^ = 0.23), while latitude explained 44% (adjusted R^2^ = 0.44). The RDA scores for latitude as well as dichromatism revealed a pattern of negative correlation with adaptive variance in southern forms of North American juncos (*phaeonotus*, *palliatus*, *dorsalis* and *caniceps*) while more boreal forms showed increasing positive association from south to north, following the phenotypic gradient of sexual dichromatism. Once again, the northernmost forms *oreganus* and *hyemalis* showed the highest positive correlation scores. In turn, *caniceps* showed low association values, especially in terms of latitude (Table 4).

**Table 4.**
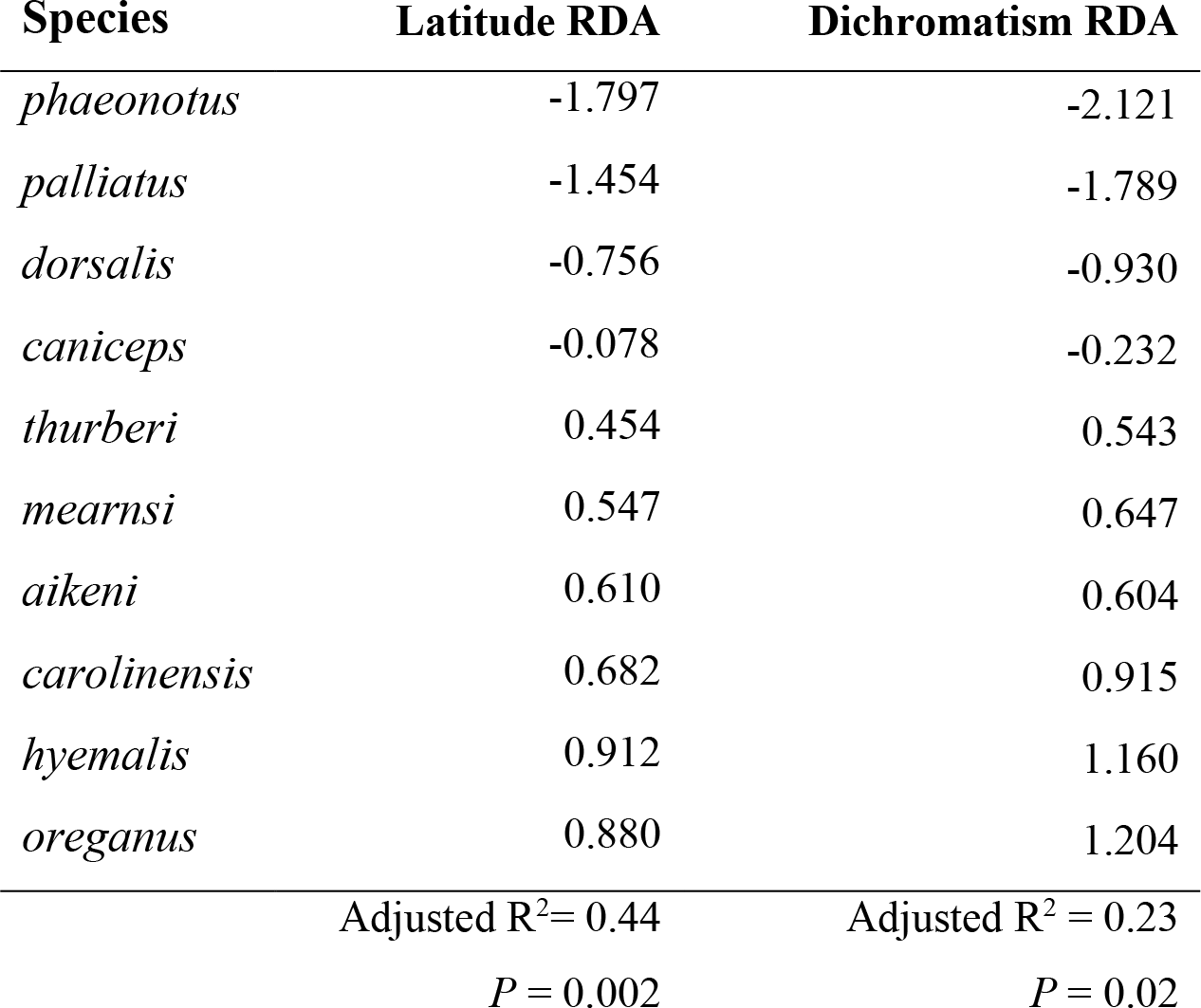
RDA scores of the constraining latitude and sexual dichromatism variables, explained variance, and *P* values. The constrained ordination tests were performed in two separate redundancy analyses, and statistical significance was computed by a permutation-based procedure with 9,999 permutations.

The variance partition analysis revealed a complete lack of orthogonality between the adaptive genetic variance explained by sexual dichromatism and that explained by latitude, i.e. the total 23% of the variance explained by sexual dichromatism was also explained by latitude, demonstrating a total overlap between their effects on adaptive genomic variance. The permutation procedure yielded a *P* value equal to 0.003, confirming the significance of the variance fraction explained by both variables. The remaining 21% of variance explained solely by latitude was also significant, with a *P* value of 0.012.

## Discussion

### Sexual dichromatism correlates with plumage coloration divergence and latitude

Our results show a strong correspondence between the strength of sexual selection and the degree of phenotypic differentiation in secondary sexual traits across the phylogenetic lineages of the genus *Junco*. Discriminant function analyses and Mahalanobis distances on colorimetric variables recovered a clear signal of plumage color differentiation for the northern, recently radiated lineages of junco, as previously reported in a similar analysis of the entire genus (Friis et al. 2016). Interestingly, the DFA of the northern lineages revealed a pattern highly congruent with the neutral genetic structure inferred in the STRUCTURE analysis, and especially in the PCA based on neutral genome-wide SNP data, congruent with the highly significant correlation between Nei’s genetic distance and Mahalanobis distances for colorimetric variables. In contrast, the DFA of morphometric variables showed low levels of clustering and high overlap among forms, as well as lower Mahalanobis distance values, suggesting weaker evolutionary pressures on ecomorphological traits than on traits potentially under sexual selection (Panhuis et al. 2001; Arnegard et al. 2010; Safran et al. 2013; Martin and Mendelson 2014).

The significant relationship between plumage color divergence and the degree of sexual dichromatism found in the simple Mantel and especially in the partial Mantel test, suggests that sexual selection may have had a major role in driving phenotypic divergence among northern junco lineages. Congruently with the higher color similarity among more closely related forms of northern junco (e.g. the yellow-eyed forms *phaeonotus* and *palliatus*; the rufous-backed forms *dorsalis* and *caniceps*; the black-hooded Oregon forms *thurberi* and *oreganus*; or the slate-colored forms *hyemalis*, *carolinensis* and *aikeni*) the correlation increased when correcting for genetic distance, supporting the existence of divergence driven by sexual selection even when comparing the most recently separated lineages. Mantel and particularly partial Mantel tests have been criticized because the permutation procedure may be an inadequate statistical significance estimator (Raufaste and Rousset 2001). However, partial Mantel tests are deemed suitable when there is low correlation between the independent variables (Castellano and Balletto 2002) as is the case in our study.

Multivariate and linear regression analyses also confirmed the increasing latitudinal pattern of sexual dichromatism from the divergent Central American lineages to the recently radiated North American forms. This signal was already proposed by Alden H. Miller in his monograph from 1941 about the genus *Junco*. Importantly, the latitudinal distribution of the *Junco* species and especially of the postglacial boreal forms reflects not only the ecological gradient across which their demographic expansion occurred, but also the historical sequence of cladogenetic events that resulted in the multiple lineages in the *Junco* complex (Friis et al. 2016). The positive correlation with latitude suggests therefore that sexual dichromatism is a derived, continuous trait that has evolved and increased during the northward recolonization and diversification of the young northern juncos, independently of the changing patterns of plumage coloration themselves.

### Interactions between sexual and natural selection

The redundancy analyses recovered signals of genetic associations for latitude and sexual dichromatism with 113 BayeScan SNP outliers, suggesting the role of both sexual selection and ecological aspects related with latitude in shaping genome-wide adaptive variability in postglacial junco forms. The ordination analyses revealed that up to 44% of the variation in adaptive variability is explained by latitude (*P* = 0.002) and consistently, the ordination scores present an association pattern that increased with latitude, with more extreme forms across the range showing the highest absolute values of correlation. Congruently with the relationship between latitude and the extent of male-female color differentiation, a similar pattern was recovered from the corresponding scores of the ordination analysis over sexual dichromatism (explained variance = 23%, *P* = 0.02). These outcomes are in contrast to the patterns obtained in the DFAs, which revealed a greater divergence in secondary sexual characters than in ecologically relevant morphometric traits among young lineages of juncos. In addition, the variance partition analysis yielded a complete overlapping between the variance explained by sexual dichromatism and latitude, showing the difficulty in distinguishing between the effects of sexual selection and latitude-dependent ecological differences.

The lack of orthogonality between latitude and dichromatism in explaining the variability on SNPs potentially under selection suggests that adaptive variation explained by sexual selection is also latitude-dependent. In other words, the fact that the total variance explained by sexual dichromatism can be explained also by latitude may indicate that adaptive variation due to sexual selection is also structured in terms of variation across the selective latitudinal axis (Lasky et al. 2012), suggesting that sexual and ecological selection may have been coupled processes in the diversification of the northern junco lineages (Butlin et al. 2012).

The association between breeding latitude and sexual dichromatism is a well-documented pattern in New World bird species, yet whether this is due to ecological factors or to non-ecological geographic variation in sexual selection remains controversial (Badyaev and Hill 2003). A similar relationship stands for migratory behavior (e.g. Friedman et al. 2009), arguably because dimorphism facilitates mate recognition and choice during shorter breeding seasons, because rapid establishment of territories increases male-male competition and intrasexual selection, or because ornamentation may be an honest signal of better performance during long seasonal movements (Hamilton 1961; Fitzpatrick 1994; but see Dunn et al. 2015). Other proposed interactions between sexual and natural selection refers to environmental constrains in the production and perception of sexual signals (Maan and Seehausen 2011) also referred as ‘external’, against ‘internal’ interactions in which ecologically adaptive traits are also sexually selected, either directly or through linked selection (Safran et al. 2013; Scordato et al. 2014). Dunn et al. (2015) recently proposed that bird coloration may be the result of the simultaneous influence of natural and sexual selection effects on two different axis, the former acting on the type of color and the latter driving male-female differences.

The overlapping signal of association of latitude and dichromatism with adaptive variance in juncos may respond to such hypothesis of migration-related adaptive advantages of sexual dichromatism, considering that migration behavior covaries with latitude. As seasonal movements increased with latitude, natural selection may have favored mate preference behavior across the different junco lineages because of the direct benefits of mating with a better quality male. Juncos present eumelanin and phaeomelanin-based plumage coloration, a type of pigment that has been shown to be an honest signal of fitness in some cases (Roulin et al. 2008; Safran et al. 2008; Maguire and Safran 2010; Scordato and Safran 2014). The gain and loss of mate preference behaviors based on such signaling to cope with the selective pressures of long distance migration is consistent with the pattern observed in the form *carolinensis*, the non-migratory subspecies of slate-colored junco from the Appalachian Mountains. In contrast to the rest of the slate-colored forms, which usually migrate long distances south of the breeding areas, the seasonal movements of *carolinensis* individuals are mainly altitudinal (Miller 1941; Nolan et al. 2002). The lesser degree of dichromatism observed in this form may reflect a relaxation of sexual selection due to sedentary habits, resulting in a reduction of male-male competition and a return to monochromatism (Fitzpatrick 1994; Badyaev and Hill 2003; Dunn et al. 2015). There is previous evidence of a reduction in sexually selected traits in juncos when shifting from migrant to sedentary habits. Yeh (2004) reported a 22% decrease in the amount of white on the tail feathers of a recently established population of *thurberi* that colonized the University of California San Diego (UCSD) campus and became year-round resident.

The amount of white in tail feathers has been demonstrated to be involved in mate choice, and to correlate with fitness traits like body size (Hill et al. 1999; McGlothlin et al. 2005), suggesting potential adaptive interactions between natural and sexual selection through honest signaling. Recently, a clear pattern of genetic structure separating UCSD residents from surrounding migratory populations and wintering individuals has been detected (Fudickar et al. 2017), which is consistent with a process of extremely fast, genome-wide differentiation driven by adaptation to a novel habitat. Another particular, intriguing case is that of the *caniceps* form. This form present moderate migratory behavior, generally moving from breeding areas in Nevada, Utah and Colorado to the mountainous sections of Arizona and New Mexico for wintering (Miller 1941). Nevertheless, the analyses recovered a signal of relatively low degree of sexual dichromatism, especially apparent in the linear regression with latitude where *caniceps* showed a high fitting deviation value. This pattern is in contrast with neighboring and closely related, less migratory forms such as *dorsalis* or southern populations of *thurberi*. While in c*arolinensis* monochromatism is seemingly a derived state lead by the loss of migratory behavior, *caniceps* represents a case of autopomorphic lack of sexual dimorphism, with no clear underlying reasons that will need further research.

The signals of genetic association recovered in our analyses are congruent with the ecological aspects and the apparent gain of discriminant mate choice behavior in the recent lineages of junco. However, there are a number of caveats and limitations in the methods applied here that need to be discussed. First, because we do not have per individual spectral and SNP data, we used population-based average values for colorimetric variables and allele frequencies, which may reduce the power of the analysis. Second, using a synthetic variable of sexual dichromatism summarizing sex differences for several distinct color variables across different plumage patches entails a simplification of its potentially complex, polygenic genetic basis. Even when this may result in a more conservative analysis, it hinders a straightforward interpretation of the inferred association signal with adaptive variance. In this sense, positive associations between allele frequencies of a reduced set of outliers and a complex synthetic variable like the sexual dichromatism index computed in this study, may be due to pleiotropic effects, variability in regulatory regions and linked variants involved in multiple color traits, common in the genetic determination of bird coloration (e.g. Poelstra et al. 2014; Toews et al. 2016; Uy et al. 2016), even across different plumage patches (Campagna et al. 2017). Third, and despite the above, the high rate of false positives (type I errors) remains a major concern in genetic association analyses. Here we followed a conservative approach by combining methods of outlier detection relying on allele frequencies (BayeScan) with association tests, aiming to reduce the rate of false positives due to factors like geographic structure, demographic history, and other distorting factors (Meirmans 2015; Rellstab et al. 2015).

### The role of sexual selection in the early stages of speciation in the *Junco* complex

There are numerous, compelling cases of rapid diversification of sexually selected traits across closely related species and populations (Price 1998; Kraaijeveld et al. 2011), both in birds (e.g. Uy and Borgia 2000; Wilkins et al. 2016; Campagna et al. 2017) and other taxonomic groups (e.g. Dominey 1984; Masta and Maddison 2002; Boul et al. 2007; Butler et al. 2007). This pattern suggests a role for sexual selection in driving phenotypic diversification at early stages of the speciation process. Several studies have also reported signs of faster evolution in sexually selected traits than in traits of ecological importance (Arnegard et al. 2010; Safran et al. 2013; Martin and Mendelson 2014), reinforcing the argument that sexual selection may contribute to diversification by increasing the rates of phenotypic change in secondary sexual traits in isolated populations (Price 1998; Panhuis et al. 2001).

The recently radiated forms of North American juncos represent one of the most striking examples of rapid phenotypic diversification, having evolved into at least six highly differentiated forms in only 18,000 years c.a. (Milá et al. 2007; Friis et al. 2016; Milá et al. 2016). These forms are not only differentiated in color and coloration patterns, but also present considerable genetic structure, suggesting that present-day contact zones among forms represent secondary contact among forms that originated in allopatry during the northward postglacial recolonization of North America. In contrast to color, minor divergence has been detected in ecomorphological traits, which supports the hypothesis of divergence arising by an increase of the overall rate of change due to sexual selection acting differentially among genetically divergent junco lineages. Under these assumptions, sexual selection driving fast phenotypic divergence may proceed independently of ecological factors (Panhuis et al. 2001; Kraaijeveld et al. 2011). However, the correlation between sexual dichromatism and latitude and the overlapping association signals of both parameters with the variability of loci putatively under divergent selection found in northern juncos is congruent with the more predominant proposed models of speciation of natural and sexual selection jointly driving diversification (Kraaijeveld et al. 2011; Maan and Seehausen 2011; Butlin et al. 2012; Wagner et al. 2012; Safran et al. 2013). Still, while sexual dichromatism is correlated with latitude, the numerous, distinct patterns of coloration of the juncos are not. At the same latitude, we can find highly divergent patterns of coloration, which is difficult to explain by ecological interactions with mate choice behavior. The stunning color diversity of juncos may be related to a process of differential mate choice in allopatric conditions: during the early stages of the postglacial recolonization process, mutations underlying color changes may have stochastically appeared in isolated populations and been positively selected through adaptive or arbitrary female choice. Due to lack of gene flow, these traits could become fixed independently among populations. High evolvability of feather color patterns and geographic isolation may thereby have fostered the rapid diversification of northern junco lineages (Schluter 2009; Nosil and Flaxman 2011; Mendelson et al. 2014). A similar hypotheses has been proposed by Winger and Bates (2015) for a number of passerine species across the arid Marañon valley of Peru, although over considerably longer periods of time.

Sexual selection may thereby promote phenotypic diversification, but the extent to which this diversification can lead to the formation of new species remains unclear (Ritchie 2007b; Kraaijeveld et al. 2011; Seddon et al. 2013). A number of studies have documented a relationship between speciation rate and the strength of sexual selection (e.g. Barraclough et al. 1995; Seddon et al. 2008; Kraaijeveld et al. 2011; Maia et al. 2013b; Seddon et al. 2013; but see Huang and Rabosky 2014) or changes in the intensity of sexual selection (Gomes et al. 2016). In recently diversified systems, sexual selection may have a predominant role as promoter of premating isolation barriers by accelerating evolutionary divergence in signals involved in species recognition, preventing admixture upon secondary contact (Price 1998, 2008). Evidence for this among northern lineages of *Junco* is mixed. Their genetic distinctiveness and highly divergent patterns of plumage coloration suggest the existence of reproductive isolation in some areas, but in others, reproductive isolation seems absent, and juncos form hybrid zones where parapatric forms come into contact. Estimates of assortative mating and hybrid fitness at these areas of introgression are lacking, and will be necessary to fully understand the degree of reproductive isolation among some junco forms. If, as hypothesized, the present parapatric limits are the result of recent secondary contact after their postglacial diversification in isolated populations, premating barriers to gene flow may have not been sufficiently developed, and the current lineages may enter in a ‘lineage fusion’ phase through extensive gene flow, erasing the incipient lineage formation (Grant and Grant 2008; Garrick et al. 2014). Alternatively, contact zones may be stable and ongoing divergence could culminate in a set of fully isolated species, which would yield a positive correlation between sexual selection strength and speciation rate at a phylogenetic level, in agreement with proposed models of speciation by means of the combined effects of sexual selection and local adaptation. In either case, the analyses reported in this study reveal a complex array of sexual and ecological factors as potential drivers of the rapid radiation of the northern lineages of *Junco*, and provide new evidence for the role of sexual selection in the early stages of lineage divergence, especially when interacting with natural selection.

## Conclusions

Our analyses confirm the ecological pattern of sexual dichromatism gradually increasing with latitude in the *Junco* system, reinforcing the hypothesis of stronger sexual selection in the North American lineages of postglacial origin. Correlation tests also demonstrated significant dependence between the degree of divergence in terms of plumage coloration and the level of sexual dichromatism, a pattern that contrasted with the lower signal of differentiation in ecomorphological traits, and suggesting that sexual selection may have been a predominant evolutionary force in driving phenotypic diversification among recently radiated forms of junco. However, redundancy analyses revealed overlapping effects of both latitude and sexual dichromatism in shaping adaptive variance, suggesting a role for sexual and ecological factors jointly driving lineage differentiation. These results, along with the patterns of neutral genetic structure of the recently radiated lineages of junco, depict a scenario of rapid divergence in isolation at early stages of the speciation process, followed a by a secondary contact phase. Whether or not barriers to reproduction have developed sufficiently to complete lineage formation, the analyses reported here reveal a complex array of sexual and ecological factors as potential drivers of the rapid radiation of the northern juncos, and provide new evidence for the proposed models of lineage divergence promoted by natural and sexual selection.

## Acknowledgements

We are grateful to the following museum curators and collection managers for allowing us access to junco specimens: Philip Unitt at the San Diego Natural History Museum (SDMNH), Kimball Garrett at Los Angeles Museum of Natural History (LAMNH), Carla Cicero at the Museum of Vertebrate Zoology (MVZ), John McCormack and James Maley at The Moore Laboratory of Zoology at Occidental College (MLZ), Chris Milensky at The National Museum of Natural History (NMNH), and Paul Sweet at The American Museum of Natural History (AMNH). We are also grateful to Rebecca Safran, Samuel M. Flaxman and Luis R. Pertierra for their kind assistance with trait divergence analyses. Funding was provided by grant CGL-2011-25866 from Spain’s Ministerio de Ciencia e Innovación to BM.

## Data Accessibility

Genomic and phenotypic data will be deposited in Dryad shortly.

## Author Contributions

GF and BM designed the study and carried out field sampling; GF generated and analyzed genomic data; GF and BM generated and analyzed phenotypic data; GF and BM wrote the manuscript.

